# Predicting metabolism during growth by osmotic cell expansion

**DOI:** 10.1101/731232

**Authors:** Sanu Shameer, José G. Vallarino, Alisdair R. Fernie, R. George Ratcliffe, Lee J Sweetlove

## Abstract

Cell expansion is a significant contributor to organ growth and is driven by the accumulation of osmolytes to increase cell turgor pressure. Metabolic modelling has the potential to provide insights into the processes that underpin osmolyte synthesis and transport, but the main computational approach for predicting metabolic network fluxes, flux balance analysis (FBA), typically uses biomass composition as the main output constraint and ignores potential changes in cell volume. Here we present GrOE-FBA (Growth by Osmotic Expansion-Flux Balance Analysis), a framework that accounts for both the metabolic and ionic contributions to the osmotica that drive cell expansion, as well as the synthesis of protein, cell wall and cell membrane components required for cell enlargement. Using GrOE-FBA, the metabolic fluxes in dividing and expanding cell were analyzed, and the energetic costs for metabolite biosynthesis and accumulation in the two scenarios were found to be surprisingly similar. The expansion phase of tomato fruit growth was also modelled using a multi-phase single optimization GrOE-FBA model and this approach gave accurate predictions of the major metabolite levels throughout fruit development as well as revealing a role for transitory starch accumulation in ensuring optimal fruit development.

## INTRODUCTION

Flux balance analysis (FBA), a method for predicting and analysing steady-state metabolic fluxes, has been widely applied in the study of unicellular and multicellular plant systems (Sweetlove and Ratcliffe, 2011; Nikoloski *et al.*, 2015; de Oliveira Dal’Molin and Nielsen, 2018). The approach requires a matrix of metabolic reaction stoichiometries and an objective function that represents the optimisation target of the biological system (Feist and Palsson, 2010). In simple unicellular organisms growing in a nutrient-rich environment, where life cycle events involve only growth and reproduction, the maximization of flux representing the accumulation of biomass elements has proved to be a reasonable objective function (Varma and Palsson, 1994; Feist *et al.*, 2007; Feist and Palsson, 2010). In complex biological systems, however, accumulation of biomass may not be the primary purpose of every cell type in the organism. For example, the principal metabolic objective of fully expanded source leaves is the biosynthesis of sucrose and amino acids for the rest of the plant (Cheung *et al.*, 2014). Moreover, organ development in plants typically involves phases of cell differentiation, cell division and cell expansion (Gonzalez *et al.*, 2012) and these are not temporally synchronous, meaning that at different stages of organ development different mechanisms of growth are dominant. Metabolism, being closely related to the demands of the cell, is thus likely to vary between these stages of development (Sweetlove and Ratcliffe, 2011; Nikoloski *et al.*, 2015). The cell expansion stage is responsible for the main increase in organ size and metabolic content in plants, and it is the dominant mechanism when growth is measured during plant phenotyping (Fahlgren *et al.*, 2015; Tardieu *et al.*, 2017). This makes understanding metabolism in expanding cells of great interest and relevance to breeding and crop engineering. In its conventional form FBA does not take account of the changing volume of the cell, indicating the need for a new FBA formulation if this objective is to be achieved.

To provide a biological context, we focused on tomato, *Solanum lycopersicum*, which has an extensive phase of cell expansion during fruit development. This process has been described in detail at the molecular-biochemical level (Valle *et al.*, 1998; Carrari and Fernie, 2006; Carrari *et al.*, 2006; Legland *et al.*, 2012; Biais *et al.*, 2014) and the available biochemical data can be used to guide the new FBA formulation. Tomato is also an important model for fleshy fruit development and ripening, owing to its agronomic value, ease of cultivation and diploid genetics. FBA has been previously used to model steady-state snapshots of tomato fruit metabolism at different stages of development (Colombié *et al.*, 2015). This was achieved by using time-series metabolomic datasets to constrain metabolite accumulation and degradation rates in a model of primary metabolism. This snapshot approach is informative, but it has three disadvantages. First, accurate predictions of metabolic state, including the accumulation of solutes during cell expansion, can only be made by imposing a large number of experimentally measured constraints. Secondly, the majority of the flux predictions are a direct consequence of the constraints imposed. For example, the rate of starch degradation observed in the model was the result of a constraint that dictates that starch is consumed at a set rate. Thirdly, because each developmental point is modelled separately in this approach, flux predictions are based on the constraints of that particular developmental point alone and the effect of future development on metabolism in earlier stages of development is lost. In our previous work on fully expanded source leaves we highlighted a similar issue with modelling day and night metabolism separately, and demonstrated that modelling day and night phases simultaneously as a single diel FBA problem led to better predictions of leaf metabolism (Cheung *et al.*, 2014). Applying a similar multiphase FBA approach to metabolism during tomato fruit development should therefore improve the usefulness of the computational approach.

In this study, we developed an innovative framework for modelling metabolism in expanding cells using FBA. This approach, GrOE-FBA (Growth by Osmotic Expansion – Flux Balance Analysis), accounts for the metabolic and ionic contributions to the osmotica that drive cell expansion, as well as the synthesis of protein, cell wall and cell membrane components required for cell enlargement. We show how GrOE-FBA can be used to identify the metabolic network fluxes in expanding cells, highlighting the major metabolic differences between dividing and expanding cells. We also show how GrOE-FBA can be combined with a multiphase single optimization FBA approach to study plant metabolism during organ development. This approach provided evidence that transitory starch stores are necessary for optimal fruit development, and that prevention of starch accumulation during the early stages of fruit development may result in smaller fruits owing to the reduced phloem uptake rate in larger tomato fruits, particularly during developmental stages where fruit expansion has been observed to be maximal.

## RESULTS

### Modelling cell expansion using GrOE-FBA

Cell expansion is driven by the accumulation of osmolytes and the resulting influx of water (Boyer *et al.*, 1985). There are two key equations (See Appendix S1 for the derivation of equations 1-5 below) that allow the osmotic content of a cell to be related to its volume, and which ultimately allow FBA to be used to model metabolism in expanding cells. First, assuming that the intracellular solutions behave as ideal solutions, the total osmotic content of a cell at steady state is equal to the product of its osmolarity (*C_cell_*) and volume (*V_cell_*)

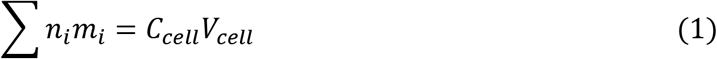

where *n_i_* and *m_i_* are the van’t Hoff factor and number of moles of metabolite *i* respectively in the cell. This equation relates changes in cell volume to changes in osmotic content, and for predicting fluxes it needs to be used in tandem with a second equation that considers the distribution of solutes between the cytoplasm and the vacuole

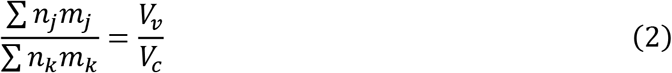

where metabolites *j* and *k* are vacuolar and cytoplasmic, respectively. This equation states that at steady state, the ratio of the osmotic content of the vacuole and cytoplasm is equal to the ratio of their volumes, *V_v_* and *V_c_* respectively.

Equations 1 and 2 provide the link between volume and metabolic content at cellular and subcellular levels, and they imply that including a representation of osmotic content in an FBA model should be sufficient to account for volume changes. This was achieved by introducing two pseudo-metabolites in the metabolic network to represent the osmoles associated with the accumulation of osmotically active metabolites/ions in the vacuole and cytosol (*O_vac_* and *O_cyt_*, respectively; Figure 1a). The pseudo-metabolites took account of the expected differences in van’t Hoff factor by constraining the model so that there could be no net change in the total charge of the vacuole or cytosol. To implement these constraints, two new pseudo-reactions, ‘aggregator reactions’, were included in the model. The first aggregator reaction, 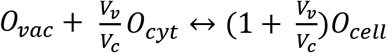, defined a pseudo-metabolite, *O_cell_*, to represent the accumulation of cellular osmolytes and constrained the model so that the ratio of accumulation of *O_vac_* and *O_cyt_* matched *V_v_*/*V_c_* (Equation 2). The second aggregator reaction drained *O_cell_* from the system with a flux equal to *C_cell_ V_cell_*. This reaction satisfied the steady-state requirement and constrained the model according to equation 1.

**Figure 1:**
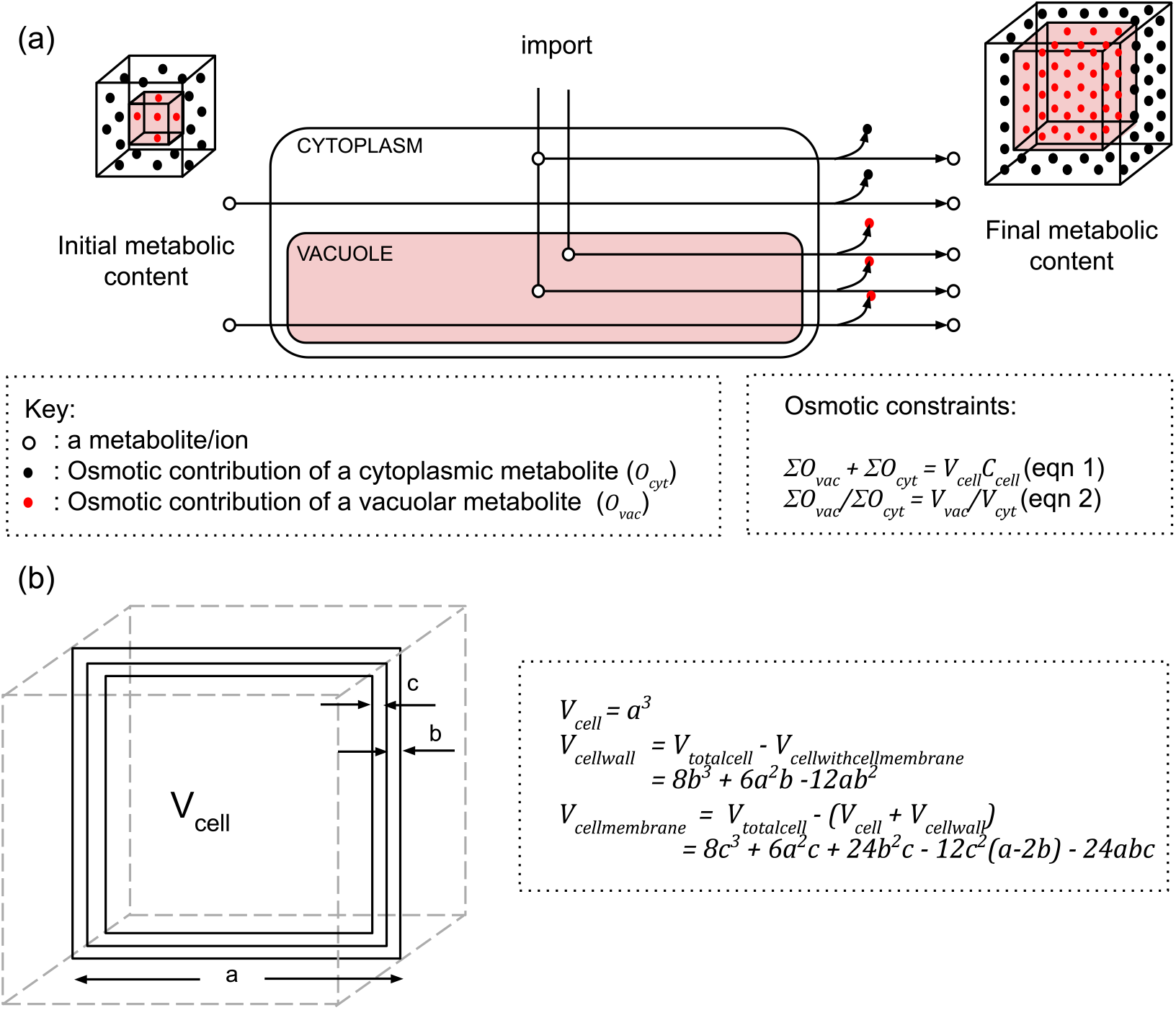
Modelling cell expansion in cube-shaped pericarp cells. (a) Cell expansion is driven by the accumulation of soluble metabolites and ions in the vacuole and cytosol. Accumulation of osmotically active species in the GrOE-FBA model is accompanied by the accumulation of pseudo-metabolites *O_vac_* or *O_cyt_*, representing the contribution of the accumulating metabolite/ion to the osmoticum of the vacuole or cytosol respectively. According to equation 1, the sum of the vacuolar and cytosolic osmoles is equal to the product of the volume (*V_cell_*) and osmolarity (*C_cell_*) of the cell. According to equation 2, the ratio of vacuolar and cytosolic osmoles is equal to the ratio of the vacuolar (*V_vac_*) and cytosolic (*V_cyt_*) cell fractions. (b) The volumes of the cell wall and cell membrane were calculated by subtracting the remaining parts of the cell from the total cell volume. *a*, edge length of the cell; *b*, cell wall thickness; *c*, cell membrane thickness. Changes in edge length lead to changes in cell volume, but the cell wall and cell membrane thickness are assumed to be constant.

As well as accumulating osmolytes, expanding cells synthesise extra cell wall, cell membrane and protein to maintain cell functions. Estimates of the additional biomass were obtained by creating a simple geometric representation of a cell assuming: (a) cells are cube-shaped; (b) the cell wall is uniformly thick and composed of cellulose only; (c) the cell membrane is uniformly thick; (d) the fraction of the total protein content of the cell in the vacuole is negligible; and (e) the cytoplasmic protein concentration is maintained during cell expansion. Based on these assumptions, if *V* is the volume of the cube-shaped cell, *b* is the thickness of the cell wall and *c* is the thickness of the cell membrane (Figure 1b), then the amount of cellulose, phospholipid and protein in the cell can be estimated from the following equations (Appendix S1)

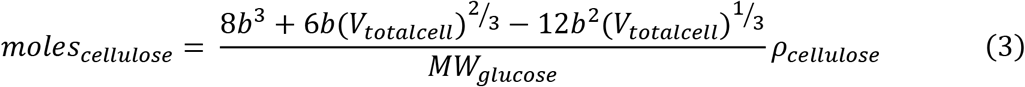

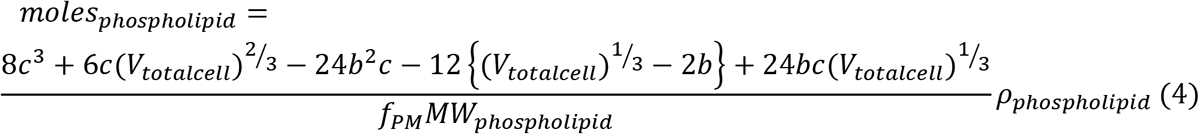

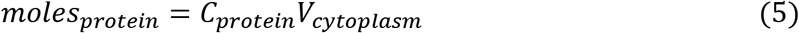

where *moles_cellulose_* is the amount of cellulose in moles, *ρ_cellulose_* is the density of cellulose, *MW_gluclose_* is the molecular weight of glucose, *moles_phospholipid_* is the amount of cell membrane phospholipids in moles, *f_PM_* is the fraction of the total lipid content found in the plasma membrane, *ρ_phospholipid_* is the density of cell membrane phospholipids, *MW_phospholipid_* is the molecular weight of membrane phospholipid, *moles_protein_* is the amount of protein in moles (based on how a unit protein is represented in metabolic models), *C_protein_* is the molar concentration of protein in the cytoplasm (based on the molar mass of a unit protein as represented in the metabolic model) and *V_cytoplasam_* is the volume of the cytoplasm.

The demand for osmolytes and biomass elements required to support a change in cell volume can be estimated from the difference in the amount of these metabolites for the initial and final cell volumes. For example, the cellulose demand for when a cell changes its volume from *V*1 to *V*2 can be calculated as follows,

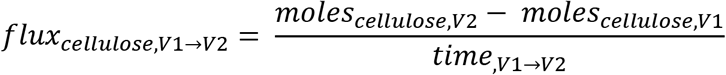

where *moles*_cellulose,*V*1_ and *moles*_*cellulose*,*V*2_ are moles of cellulose when the cell has a volume of *V*1 and *V*2, respectively; and *time*,_*V*1→*V*2_ is the time the cell takes to change its volume from *V*1 to *V*2.

In this manner, by combining previously described osmotic constraints (based on equations 1 and 2) with biomass constraints (based on equations 3, 4 and 5), it is possible to perform FBA while accounting for cell volume.

### Validation of the equations used to estimate the change in biomass during cell expansion

The cell wall, lipid and protein contents of tomato pericarp cells during fruit development were determined (Data S1) and the results compared with predictions based on equations 3-5 (Figure 2). Pericarp cell membrane phospholipid composition was assumed to be composed of phosphatidyl ethanolamine (PE), phosphatidyl choline (PC) and phosphatidic acid (PA) based on published data in cherry tomato (Guclu *et al.*, 1989) and the relative amounts of PE, PC and PA were used to estimate values for *ρ_phospholipid_* and *MW_phospholipid_*. The value of *f_PM_* was estimated from data published in tobacco leaves (Cacas *et al.*, 2016) and assumed to be constant throughout fruit development. Values of other parameters required in the equations were collected from published data or measured experimentally (Table S1).

**Figure 2:**
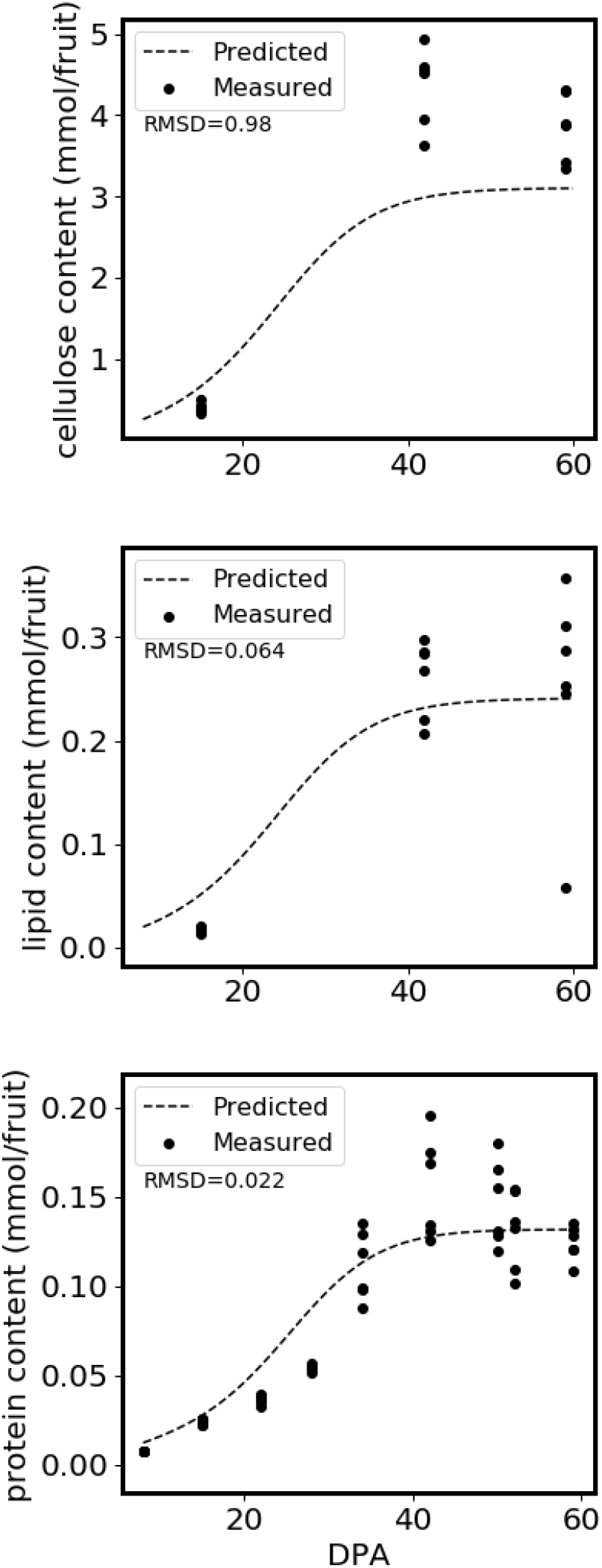
Cellulose, lipid and protein contents of tomato pericarp cells. The predicted curves are based on equations 3-5 for cellulose, membrane phospholipid and protein content respectively.

Figure 2 shows that equations 3-5 generated a good match to experimentally measured values for tomato fruit. The slight underestimation of fruit cellulose content is likely due to the assumption of the cells being geometric cubes with a constant cell wall thickness, which does not account for increased cell wall thickness around corners of the cell (Legland *et al.*, 2012). Thus, a simple geometric model can be used to predict increases in cellulose, phospholipid and protein content during cell expansion, and this allows these major biomass outflows from the metabolic network to be estimated, avoiding the need for direct measurements.

### Application of GrOE-FBA to expanding tomato cells

Because of the energy cost of synthesising cellular macromolecules such as protein, lipid and carbohydrates (Schwender and Hay, 2012), it is generally assumed that synthesis of new cells is more expensive than expanding existing cells (Lynch, 2019; Taiz, 1992). However, even during cell expansion, new macromolecules need to be produced and the synthesis and accumulation of osmolytes to drive cell expansion also represents a significant cost. The GrOE-FBA framework allows the costs of cell expansion to be accurately computed and can be compared to a computation of the costs of cell division computed using conventional FBA with a biomass objective. To do this, we modelled rapidly dividing tomato pericarp cells in culture using data from (Rontein *et al.*, 2002) and compared the resulting model to one of rapidly expanding pericarp cells in fruit. Both models were constructed using a minor updated version (PlantCoreMetabolism_v1_2) of our previously published core stoichiometric model of primary plant metabolism (Shameer *et al.*, 2018). To make the models comparable, they were provided with the same precursors: glucose as the sole carbon source, as well as NH_4_^+^, O_2_, PO_4_^-^ and SO_4_^-^. The same objective function of minimization of the sum of fluxes was applied to both models. Experimental data was used to constrain the rate of biomass production in the dividing cell model to 2 mg DW/mL/day, the fastest rate reported in the Rontein et al. (2002) study. The rate of cell expansion in the expanding cell model was set to that of fruit at 26 days post anthesis (DPA) where the fastest rate of fruit growth was observed (Beauvoit *et al.,* 2014). The initial exploration of the expanding cell model included an additional component to the objective function: to maximise the organic solute content while satisfying the osmotic constraint (see subsequent explanations and discussions).

Figure 3 summarizes the results of this comparison for equivalent numbers of cells and the complete list of predicted fluxes is provided in Data S2. Comparison of the metabolic fluxes in the two models revealed that dividing cells have higher fluxes of cellulose, protein, starch and lipid biosynthesis as expected; while the expanding cells accumulated significantly more organic solutes, both in the vacuole and cytosol (Figure 3a and 3b). This accumulation of organic solutes in the expanding cell model increased the carbon demand of the model, and somewhat surprisingly, the overall rate of glucose consumption was substantially higher in the expanding cell model (5.82 mg/mL in a day) than in the dividing cell model (3.57 mg/mL/day). The flux maps (Figure 3 a,b) reveal that a substantial proportion of this additional glucose (66% of carbon taken up by the cell) was transported into the vacuole to satisfy the osmotic requirement for cell expansion. But it is also apparent that there were comparably high fluxes of glycolysis, and even higher TCA cycle flux, relative to the dividing cell model, suggesting that the energy demand of cell expansion is comparable to that of cell division. To provide a more precise quantitative comparison of energy costs, we constructed an energy budget in the two systems by collating all the fluxes in the models that required ATP consumption or led to ATP production (Figure 3d). The total ATP demand of the expanding cells was 0.16 mmol ATP/mL/day (0.10 mmol ATP/mL/day when excluding non-growth associated maintenance or NGAM) and that of the dividing cells was 0.14 mmol/ATP/day (0.08 mmol ATP/mL/day when excluding NGAM). This showed that indeed both models have very similar energy demands. Looking at the breakdown of energy expenditure (Figure 3 d) it is apparent that although the expanding cell model devoted considerably less ATP to synthesis of protein, lipid and cell wall (12 % compared to 44% in dividing cells), this was offset by a large increase in the cost of solute biosynthesis and accumulation which represented 50% of the ATP budget in the expanding cell model compared to only 11% in dividing cells (Figure 3d).

**Figure 3:**
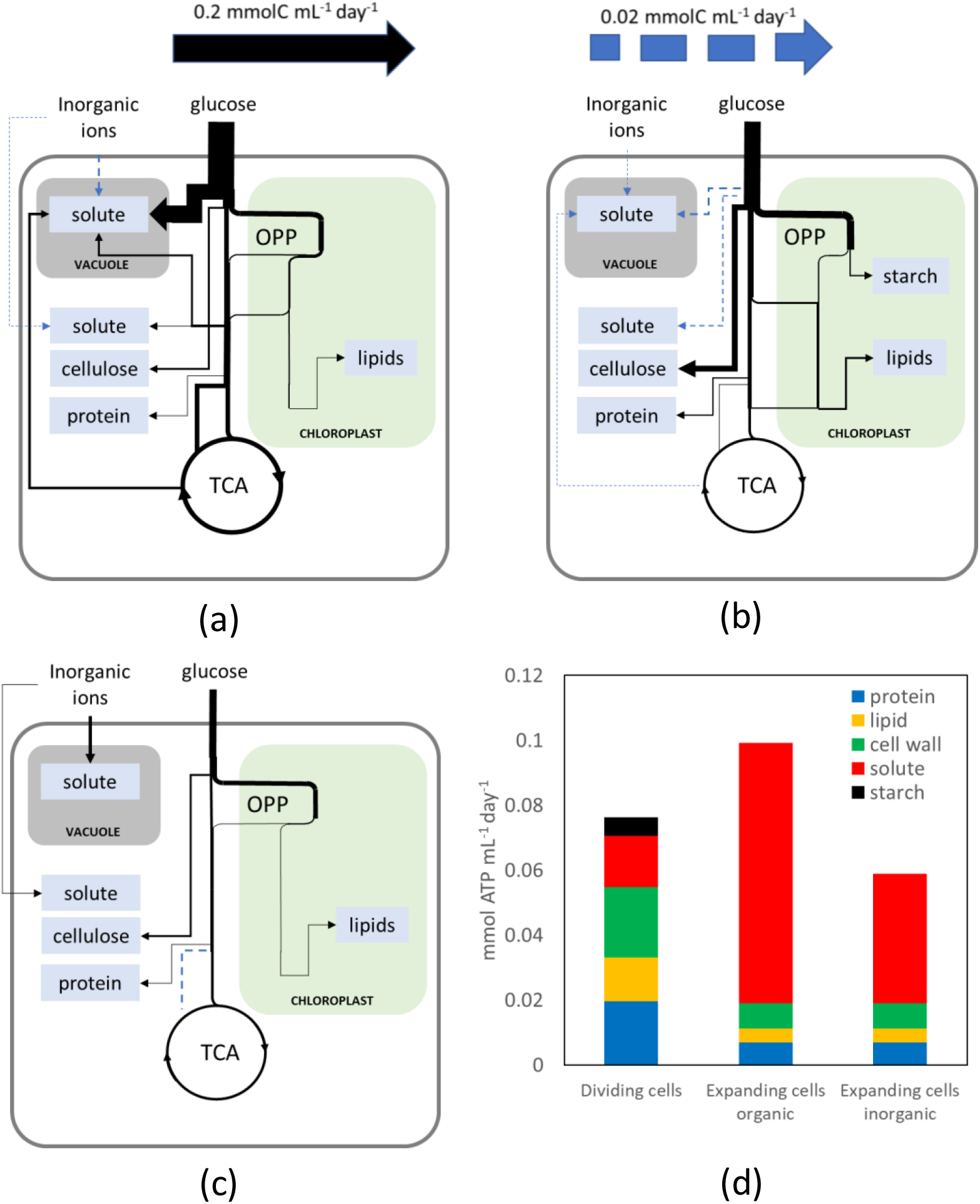
Flux prediction in expanding and dividing tomato cells using GrOE-FBA. Predicted metabolic fluxes in: (a) expanding cells accumulating only organic solutes; (b) dividing cells; and (c) expanding cells accumulating only inorganic solutes. Organic and inorganic fluxes are represented as mmolC/mL/day and mmol/mL/day respectively with the thickness of the lines scaled to match the fluxes. Values of all the predicted fluxes are given in Data S2. (d) ATP budgets for solute accumulation and for the biosynthesis of protein, lipid, cell wall, and starch, deduced from the three predicted flux maps.

Given that maximization of organic solute was a component of the objective function of the expanding cell model, the very high rate of accumulation of glucose to satisfy the osmotic constraint may be a feature of how we set up the model. Although it is known that fruit cells do contain high amounts of hexose sugars (Biais *et al.*, 2014), it is likely that inorganic solutes contribute to the osmotic balance and this would affect the total energy cost of osmolyte production and accumulation. To explore this, we removed the maximization of organic solute objective and just used minimization of the sum of fluxes as the sole objective, as in the dividing cell model. This led to a significantly lower glucose consumption rate in expanding cells (1.61 mg/mL/day) but the overall ATP demand remained comparable to the previous model (0.12 mmol ATP/mL/day). The predicted flux distribution (Figure 3c) revealed that the model was exclusively using inorganic ions to satisfy the osmotic constraints. In reality expanding cells are expected to accumulate both organic and inorganic solutes to facilitate cell expansion, so the predicted energy demand of the expanding and dividing cells are comparable. These results demonstrate that cell expansion is not necessarily a cheap option for plant growth as previously argued (Taiz, 1992).

Many similarities between the three systems (dividing cells, expanding cells accumulating organic solute and expanding cells accumulating solely inorganic solute) were apparent. For example, in each system, more than 75% of the ATP demand was met by mitochondrial ATP synthesis (75%, 76% and 79% respectively) and NGAM was found to be a significant energetic drain (45%, 39% and 51% of the total ATP demands respectively). The second and third largest energetic demands were the ATP cost of maintaining the plasma membrane proton gradient (PM-ATPase) during nutrient uptake (18%, 21% and 30% of the total ATP demands respectively) and the hexokinase flux (14%, 14% and 7%). While the basis for NGAM was identical in the three models, the PM-ATPase and hexokinase flux are dependent on the carbon demands of the model. As a result, the energy demand of the system is significantly influenced by the carbon demand for biosynthesis.

### A multi-phase metabolic model of primary metabolism in developing tomato fruit

To investigate the changing metabolic requirements for growth by cell expansion during organ development, pericarp metabolism was modelled using GrOE-FBA in the expansion and ripening stages of developing tomato fruit by dividing the time-course from 8 days post anthesis (DPA) to red ripe stage (59 DPA) into ten developmental substages. Ten copies of PlantCoreMetabolism_v1_2, each representing one of these substages, were combined to generate a 10-phase constraint-based model of primary metabolism in tomato pericarp cells. To allow for the accumulation and utilisation of sugars (glucose, fructose and sucrose), amino acids (including GABA) and organic acids (malate and citrate) in each phase, reactions transferring metabolites and ions from one phase to the next, called “linker reactions”, were added to the cytosol and vacuole of the model. Similarly, transfer of starch from one phase to the next (allowing for accumulation and utilisation of starch) was enabled via linker reactions in the plastid. The linker reactions and the multiphase nature of the model ensure that metabolism in each stage of fruit development depends not only on metabolism in the previous developmental stages but is also influenced by that of future developmental stages. Each phase in the multiphase also had access to the xylem and phloem, and was subjected to biomass requirements. A full schematic representation of the model is presented in Figure 4.

**Figure 4:**
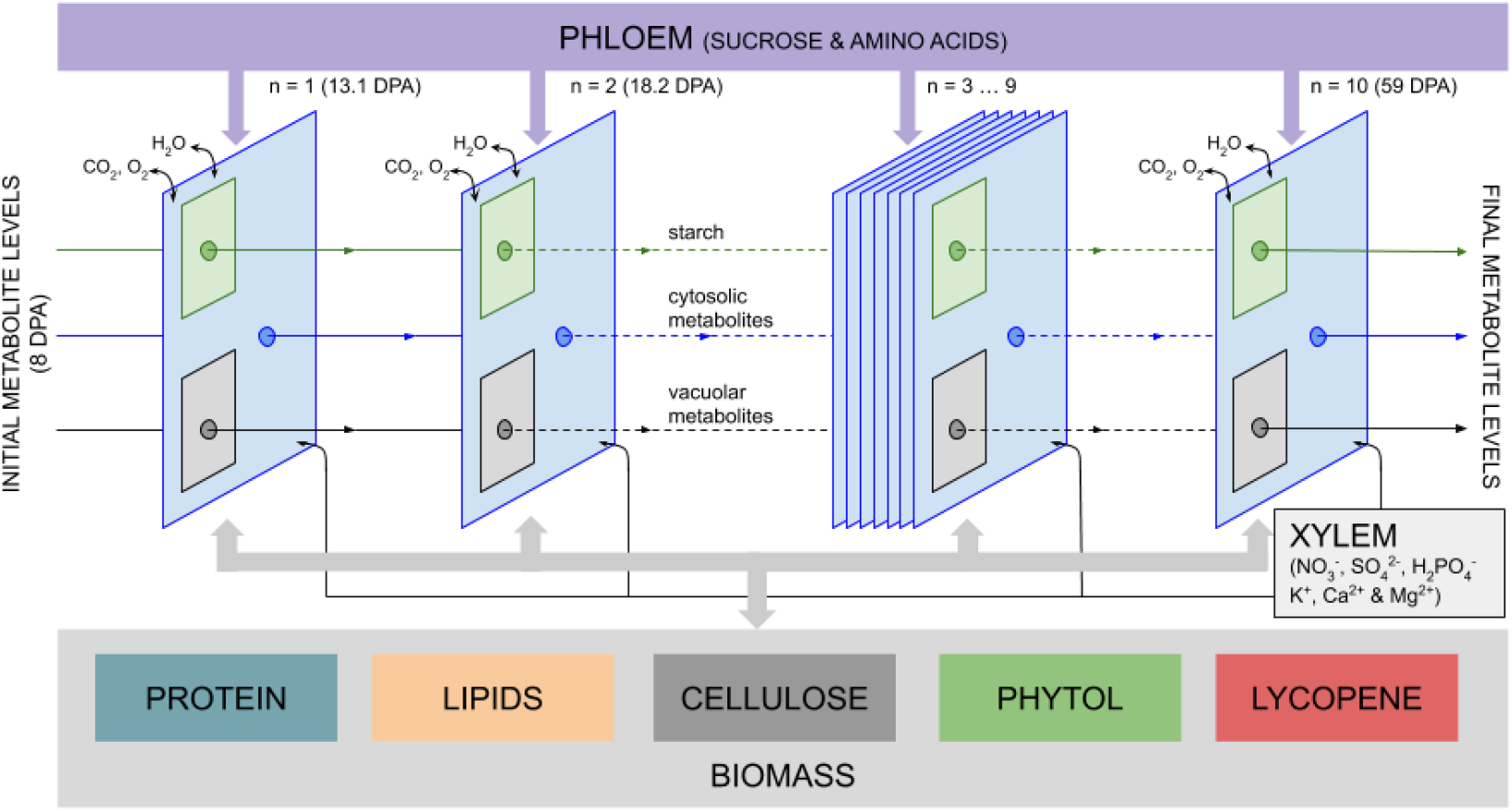
Schematic representation of the multiphase model of tomato fruit development. Fruit development from 8 days post anthesis (DPA) to 59 DPA was divided into 10 phases. Each phase has access to the indicated nutrients from the mother plant via xylem and phloem and is subject to biomass demands. Accumulation of certain metabolites and ions were permitted in each phase, facilitated in the model by ‘linker reactions’ that allow the accumulated metabolites / ions to be passed to the next phase.

The initial metabolite content of the fruit pericarp at 8DPA was established by adding source reactions (reactions that represent the import of metabolites from outside the modelled system) for sucrose, fructose, glucose, amino acids, GABA, organic acids and starch to the first phase of the model. The flux through these source reactions was constrained using the experimentally determined soluble metabolite content of 8 DPA fruits (Data S1). The soluble metabolite content of red ripe tomato fruits (59 DPA) (Data S1) was used to constrain the relative proportions of the soluble metabolites in the final model phase of the model by introducing appropriately constrained sink reactions. Previously published data on vacuolar pH during fruit development in cherry tomatoes (Rolin *et al.*, 2000) was used to set the vacuolar pH for the ten phases and thus the abundance of the different metabolite charge states for each phase. A data-based constraint was also applied for the maximal rate of uptake of nutrients into the fruit from the phloem (see Materials and Methods). The multiphase developing fruit model was solved as a single parsimonious FBA (pFBA) problem, an approach similar to previously published multiphase (Cheung *et al.*, 2014; de Oliveira Dal’Molin *et al.*, 2015; Shameer *et al.*, 2018) and multi-tissue (Grafahrend-Belau *et al.*, 2013; de Oliveira Dal’Molin *et al.*, 2015; Scheunemann *et al.*, 2018; Shaw and Cheung, 2018) constraint-based models, with the maximization of fruit soluble metabolite content as the objective. Fruit content and composition is key in increasing yield in tomato cultivation, hence, the maximization of flux through the reaction representing the accumulation of soluble metabolites in ripe tomatoes was deemed an apt objective function for the model. NGAM, nutrient uptake from the phloem and the osmotic constraints based on equations 1 and 2 were implemented as described in Materials and Methods.

### A multiphase developing tomato fruit model predicts realistic final fruit metabolite content

Figure 5 depicts the influence of the various constraints on the predictions of the multiphase developing fruit model. Optimal flux distributions of the model without the osmotic and biomass constraints to describe cell expansion and with no upper limit on phloem uptake rates predicted unrealistically high fruit organic solute content (~5000 mOmol/fruit; Fig. 5a). The introduction of osmotic constraints on the system greatly reduced the final fruit organic solute content to 24.5 mOsmol/fruit (Fig. 5b). This value was just 7.5% lower than the measured value (26.5 mOsmol/fruit). The imposition of biomass demand fluxes did not affect the predicted ripe fruit organic content, although it did cause small variations in predicted fruit content during the early stages of fruit expansion (Fig. 5c). Limiting the phloem uptake rate had no effect on the predicted organic content of ripe fruits, but it did reveal a requirement for a transitory carbohydrate store in expanding fruits (Fig. 5d).

**Figure 5:**
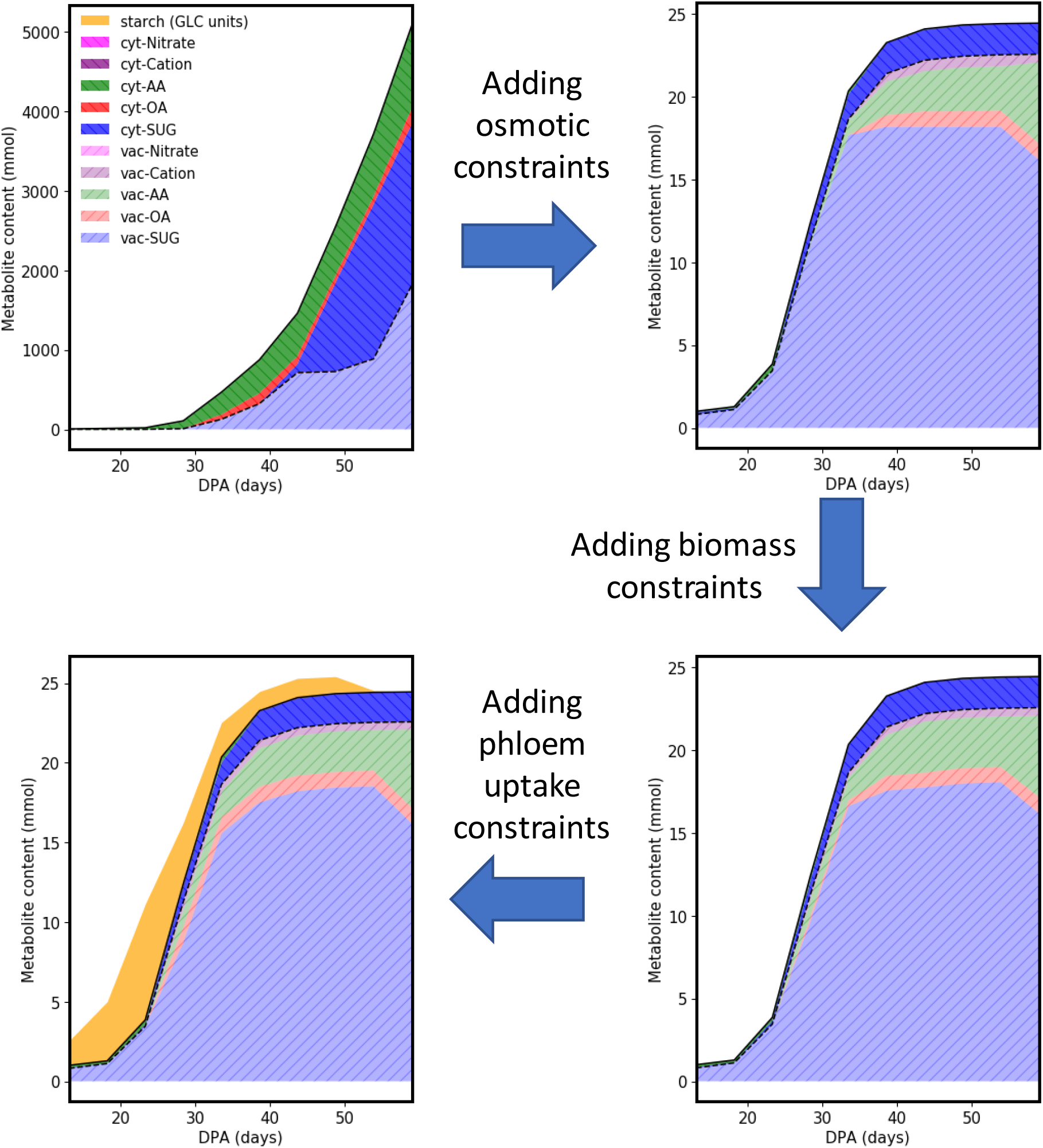
Impact of model constraints on the predicted metabolite contents of developing tomato fruit. The charts show the sequential effect on the predicted metabolite contents of adding osmotic, biomass and phloem uptake constraints to the model. In the absence of these constraints the predicted total metabolite content greatly exceeded the expected value, whereas after applying the constraints there was close agreement between the measured and predicted values. AA, amino acids; OA, organic acids; SUG, sugars (glucose, fructose and sucrose).

In order to understand the factors driving metabolism during fruit development, predicted fruit metabolic content during development was analysed (Table 1). From Table 1, it can be seen that the soluble organic content contributes a significant fraction of fruit content particularly during the later stages of fruit development. On the other hand, cellulose, lipid and protein content form a fixed fraction of predicted fruit content throughout development. The optimal flux distribution and a fully constrained version of the multiphase model are available in Data S2 and Data S3, respectively.

**Table 1:** Predicted increase/decrease in fruit content by the multiphase developing fruit model.

| Phase<br>(DPA) | fruit content (% fruit DW) |  |  |  |  | fruit DW<br>(mg) |
| --- | --- | --- | --- | --- | --- | --- |
|  | Cellulose | Phospholipids | Protein | Starch | Organic solutes |  |
| 8-13.1 | 9.49 % | 2.81 % | 1.1 % | 27.6 % | 59.0 % | 927.6 |
| 13.1 – 18.2 | 8.04 % | 2.39 % | 0.91 % | 34.8 % | 53.9 % | 1989 |
| 18.2 – 23.3 | 6.35 % | 1.88 % | 0.72 % | 31.0 % | 60.1 % | 4104 |
| 23.3 – 28.4 | 7.45 % | 2.21 % | 0.87 % | 13.4 % | 76.1 % | 4811 |
| 28.4 – 33.5 | 8.21 % | 2.44 % | 0.97 % | 6.20 % | 82.2 % | 5207 |
| 33.5 – 38.6 | 8.33 % | 2.47 % | 0.99 % | 2.92 % | 85.3 % | 5587 |
| 38.6 – 43.7 | 8.15 % | 2.42 % | 0.97 % | 2.74 % | 85.7 % | 5930 |
| 43.7 – 48.9 | 8.33 % | 2.47 % | 0.99 % | 2.35 % | 85.9 % | 5869 |
| 48.9 – 54.0 | 8.78 % | 2.61 % | 1.04 % | 0.27 % | 87.3 % | 5562 |

### Comparing model predictions to experimental measurements

Figure 6 compares the fruit content of selected metabolites predicted by GrOE-FBA with the experimentally determined values. The complete dataset for 22 solutes is provided in Supplementary Figure S1. The model was capable of making accurate predictions for glucose, fructose, glutamine and glutamate, despite being free to choose which metabolites it accumulated to satisfy the osmotic constraint. These four metabolites formed 88.6 % of organic osmolytes in the pericarp cells and their accurate prediction by the osmotically-constrained model suggests that their pattern of accumulation in developing tomato fruit represents an efficient way to drive osmotic expansion. Some other metabolite levels, including those of malate and citrate, were predicted with reasonable accuracy during the later stages of fruit development; while for others, including sucrose and GABA, it was not possible to predict the observed pattern of accumulation. This suggests that the accumulation of these metabolites is not primarily osmotically driven and that other aspects of fruit physiology may drive their accumulation during fruit development. For example, factors such as insect resistance, texture and flavour, all have an impact on determining fruit composition during development (Tohge *et al.*, 2014; Cohen *et al.*, 2014; Takayama and Ezura, 2015). Translating these factors into constraints for a metabolic model would be challenging, since the measured content of some of the poorly-predicted amino acids (aspartate and serine) fell within the FVA ranges of the model (Figure S1) but applying extra constraints to the system may improve model prediction.

**Figure 6:**
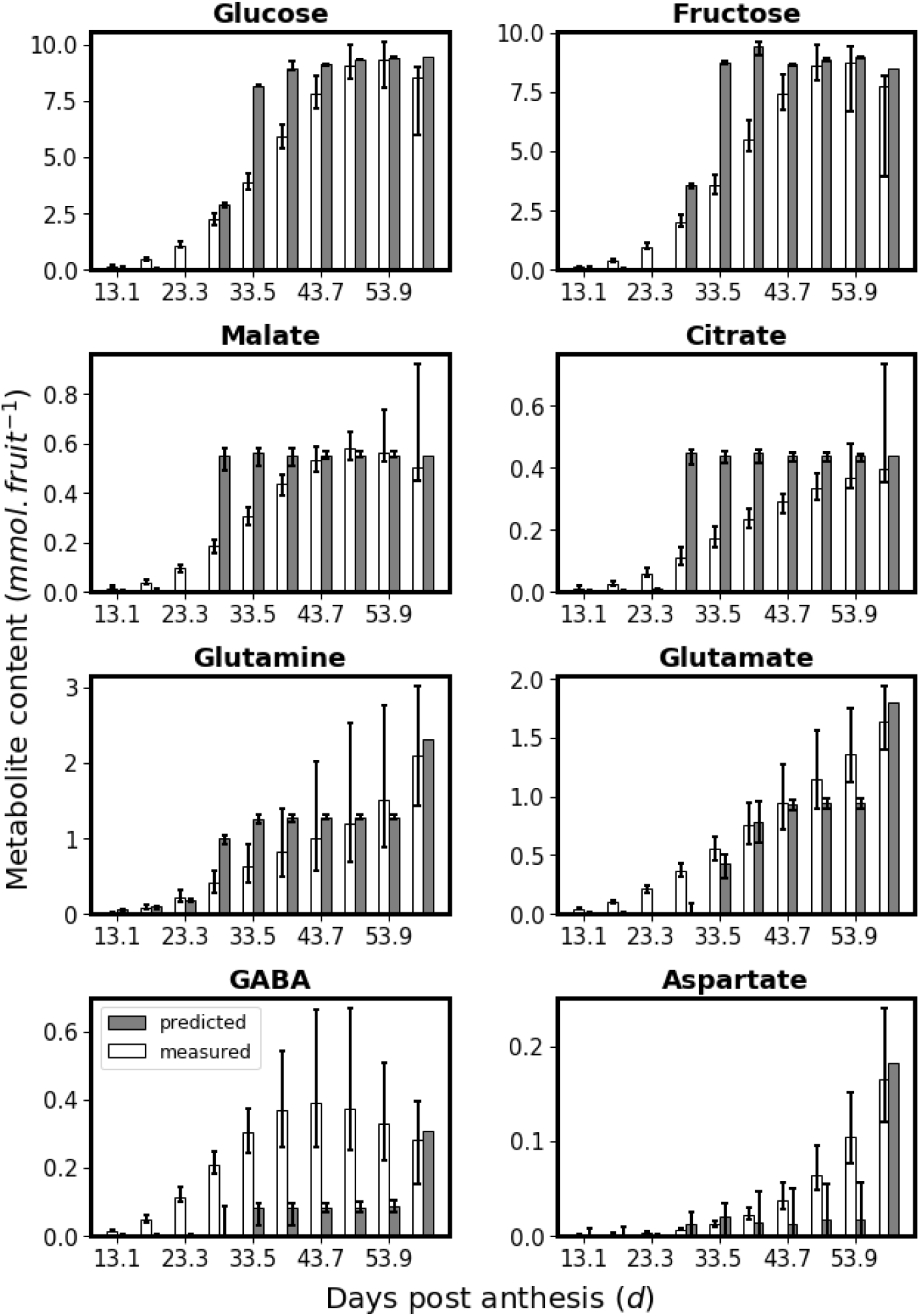
Comparison of predicted and measured metabolite contents in developing tomato fruit. The predicted metabolite contents are in good agreement (glucose, fructose, glutamine, glutamate), partial agreement (malate, citrate) or no agreement (GABSA, aspartate) with the measured values. The error bars correspond to the FVA ranges of the predicted values.

### A transitory carbohydrate store is required when phloem nutrient uptake is limited

If nutrient influx from the phloem was unlimited, then the nutrient requirement for fruit development was predicted to peak in phases 4 and phase 5 (23.3 – 33.5 DPA; Fig. 7a). In contrast when an upper bound on phloem uptake (Fig. 7b) was imposed, based on experimental data, the model predicted the need for transitory carbohydrate storage (Fig. 7c), a well-documented behaviour of tomato fruits (Ho and Hewitt, 1986; Gillaspy *et al.*, 1993). When phloem uptake rate limits were removed from the model there was no accumulation of starch (Figure 7d), suggesting that the accumulation of starch was associated with the phloem influx limits and metabolic demand. Moreover, preventing the accumulation of starch caused a 40% drop in the organic content of the fruit, demonstrating the significance of transitory starch storage in developing tomato fruits.

**Figure 7:**
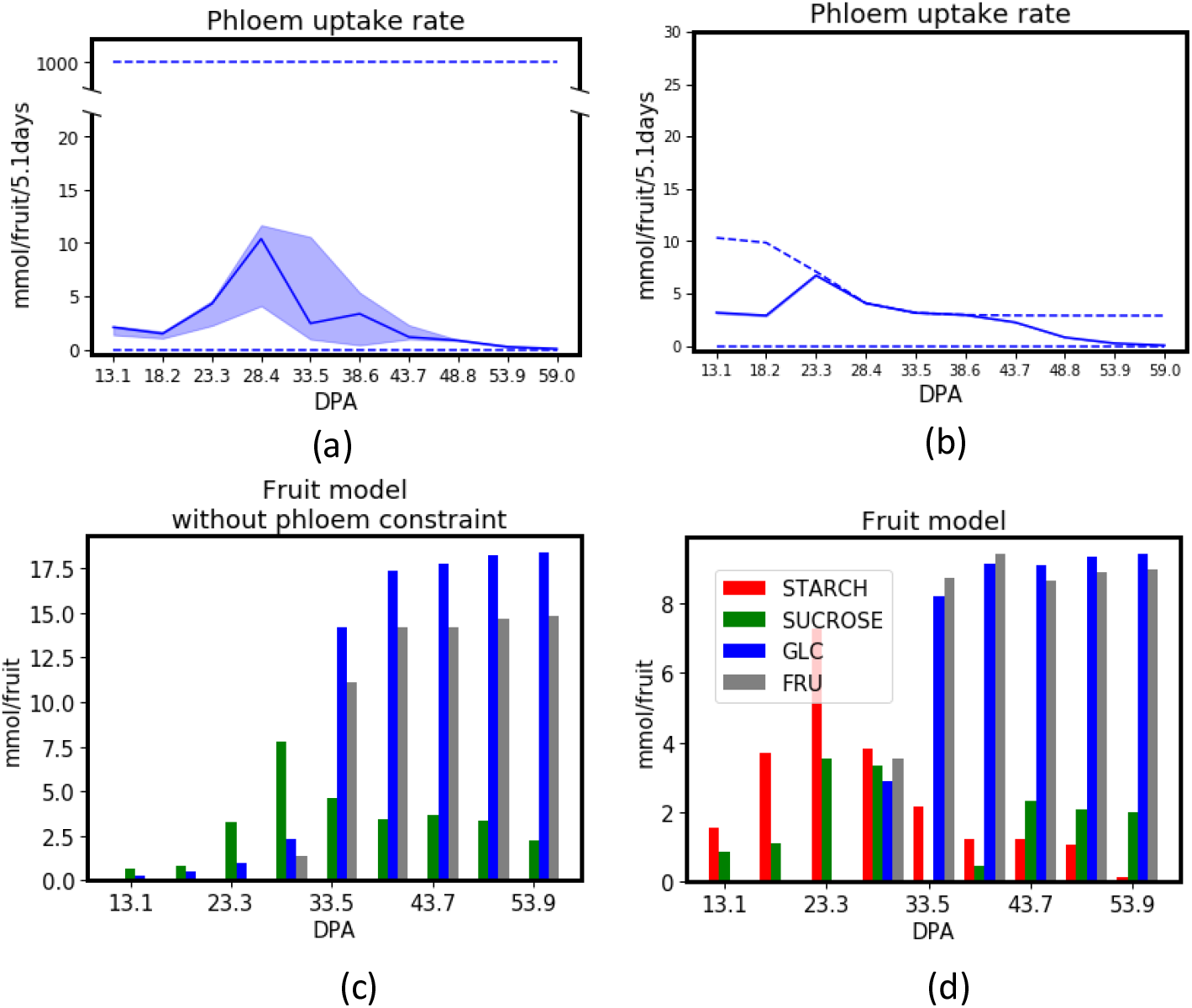
Impact of phloem uptake rate on the need for a transitory carbohydrate store during tomato fruit development. The multiphase GrOE-FBA model was run with the phloem uptake rate either (a) effectively unconstrained (allowing the model to choose a rate between 0 and 1000 mmol/fruit/5.1 days) or (b) constrained with the upper bound based on the measurements of Walker and Ho (1977a,b). The upper and lower bounds of the phloem uptake rate are shown as dotted lines, and the optimal value chosen by the model as a solid line, together with the FVA range (shaded) for the solution. Applying the upper bound to the phloem uptake rate creates a carbon and/or energy limitation in phases 4 and 5 (28.4 – 33.5 DPA) of the multiphase fruit model. The predicted starch and soluble sugar contents obtained with the phloem uptake rate either (c) unconstrained, or (d) constrained, show that phloem uptake rate constraints are responsible for the temporary carbohydrate buildup observed in the early stages of fruit development.

## DISCUSSION

### GrOE-FBA: proof of concept in the comparison of expanding and dividing cells

GrOE-FBA was developed to predict metabolic network fluxes in cells undergoing osmotically-driven cell expansion. As a proof of concept, GrOE-FBA was used to highlight the differences between metabolism in dividing and expanding cells, and the analysis led to the conclusion that cell expansion does not necessarily put a smaller requirement on cellular resources than cell division. While it is not surprising that differences existed between metabolism in dividing and expanding cells, quantitative network-level comparisons have not been made before. The two systems chosen here for comparison were dividing cells in heterotrophic cell suspension culturesand expanding cells in developing tomato fruits. A more appropriate comparison would be to model metabolism during the very early phase of tomato fruit development, when cell division dominates, and the later phases when expansion dominates. However, there is very little biochemical data to constrain such a cell division model, due to the very small size of young plant organs in the cell division phase. Even so, the marked difference in the predicted metabolic fluxes of dividing and expanding cells (Fig. 3) implies that FBA studies on plant tissues such as growing leaves (Yuan *et al.*, 2016; Shaw and Cheung, 2018) and roots (Grafahrend-Belau *et al.*, 2013; Shaw and Cheung, 2018), which generally use a biomass objective even though measurable growth in such organs is largely by expansion, could be predicting erroneous flux distributions.

### GrOE-FBA: application to tomato fruit development

GrOE-FBA was combined with multiphase modelling to analyse metabolism in developing tomato fruit. The approach required minimal constraints: a knowledge of the cell volume changes during expansion, an estimate of the volume ratio of vacuole to cytosol, and an estimate of protein concentration (*C_protein_*, Eq. 5). Using these constraints, it was possible to make surprisingly accurate predictions about the accumulation of biomass components and the principal osmotic metabolites that drive cell expansion in tomato fruit (Figs 5 and 6). The same approach could be applied to the growth of other organs such as leaves and roots.

Previous modelling studies taking into account cell expansion and metabolism were based on analysis of enzyme kinetics and were capable of exploring only limited aspects of metabolism because of the difficulty of gathering or estimating large numbers of kinetic parameters (Beauvoit *et al.*, 2014). Previous work using FBA modelling to analyse metabolism in developing tomato fruits used separate models for each stage of development (Colombié *et al.*, 2015). Such highly constrained models can provide answers to the question “What is happening?” but cannot answer the question “Why is it happening?”. For example, with this approach it has been shown that breakdown of transitory starch is the underlying cause of the respiratory climacteric in tomato fruit (Colombié *et al.*, 2017) but it was not possible to explain the purpose of the transitory starch accumulation. This is because starch accumulation and degradation were constraints of the model and not predictions. In contrast, starch accumulation and degradation in the multiphase GrOE-FBA model are system-level predictions that depend on the metabolic demand on the system and the availability of nutrients from the phloem, throughout development.

### GrOE-FBA: insights into tomato fruit development

Two major biological insights emerged from the analysis of the GrOE-FBA model of tomato fruit development. First, we demonstrate that transitory carbohydrate stores are only predicted by the model if a constraint is added that gradually decreases the rate of influx of nutrients from the phloem. Our model demonstrates that due to the decline in phloem influx during development, accumulation of starch (or other carbohydrates such as sucrose) in the earlier stages of fruit development is required to meet the carbon demand in later stages. The question then arises as to why the phloem influx into a fruit declines below that which is required to maintain its metabolic requirements. One possibility is that due to the staggered initiation of fruits along a truss, it is necessary to progressively reduce the share of phloem nutrients taken by maturing fruit to allow younger fruits, that have lower sink strength, to develop successfully. Temporal extension of starch synthesis has been reported to result in increased fruit size (Petreikov *et al.*, 2006; Petreikov *et al.*, 2009). This is in agreement with our hypothesis. Increased sucrose demand owing to starch synthesis will increase the sink strength of smaller fruits. Once the fruits are bigger, the greater transitory starch stores will reduce their sink strength, resulting in more phloem constituents being available to the smaller fruits.

The second biological insight is the relatively minor impact biomass demand had on developing fruit metabolism (Fig. 4). It is known that cell expansion results in an increase in cell wall and cell membrane content. Protein content of cells is also thought to increase in order to maintain the optimal concentration required for unhampered metabolism. In most FBA models of growing plant tissues to date, synthesis of these biomass components is the main drain on the metabolic system. The GrOE-FBA model demonstrates that for tissues growing by expansion of existing cells, demand for these biomass elements imposes only a small fraction of the metabolic cost. Instead, the dominant demand on metabolism was solute accumulation for osmotically-driven cell expansion. Accumulation of solutes was responsible for most of the gain in fruit DW during 18-43.7 DPA (Table 1). From this we can conclude that the drivers for metabolism in expanding cells are significantly different from dividing cells and that osmolarity-based constraints such as GrOE-FBA applied in this study, are necessary to model metabolism in tissues growing by cell expansion.

### GrOE-FBA: limitations of the approach

GrOE-FBA is a modified form of FBA and hence inherits its limitations (Sweetlove and Ratcliffe, 2011), including not being able to predict futile cycles. A substantial fraction of the energy consumption in a tomato cell suspension has been attributed to the operation of such cycles (Rontein *et al.*, 2002) and the FBA model is unable to capture this energy expenditure. While this might explain the discrepancy between the glucose consumption rate of the model (3.57 mg/mL/day) and the value reported in the literature (glucose influx rate for day 4 is 5.34 mg/mL/day), it should also be noted that the method used to estimate the futile cycles has been shown to be confounded by the subcellular structure of the metabolic network (Kruger *et al.*, 2007).

Another limitation of GrOE-FBA, is that it focuses on the osmotic significance of the accumulating solutes and makes no attempt to tackle the challenges involved in imposing constraints on metabolism related to insect resistance, texture and flavour, Nevertheless, introducing such constraints into the model might improve its ability to predict sucrose and organic acid levels.

Finally, the GrOE-FBA model predicts metabolic fluxes by flux optimization and does not account for signalling and regulation of metabolism, although these could be included in the form of experimental constraints on maximal enzyme activity. As a result, the model we present does not activate flux through energy dissipation mechanisms (such as alternative NADH dehydrogenases and alternative oxidase) and hence the GrOE-FBA model is unable to predict the respiratory climacteric (Colombié *et al.*, 2017). A computationally expensive FBA approach called cost-weighted flux minimization has been shown to be capable of predicting alternative pathways in source leaves (Cheung *et al.*, 2015). Such an approach might allow the multiphase GrOE-FBA tomato model to predict metabolically inefficient processes such as AOX flux and futile cycles.

## CONCLUSION

GrOE-FBA, a novel FBA approach that uses volume-based osmotic constraints allows metabolism to be modelled during cell expansion, and thus extends the scope of constraints-based metabolic modelling to growing tissues in which expansion of existing cells is the dominant driver of growth. Understanding metabolism in expanding cells could help develop engineering strategies to optimize metabolism during the cell expansion phase of leaf and root development. The approach could also be used to study metabolism in guard cells where turgor-pressure-driven changes in cell size regulate stomatal aperture, in turn regulating CO2 assimilation and transpiration rates. GrOE-FBA could therefore help complement current kinetic models of guard cells (Hills *et al.*, 2012) with a more holistic modelling approach. Recently, there has been increased interest in combining metabolic models with whole plant developmental models (Marshall-Colon *et al.*, 2017). Growth of sink tissues in the metabolic component of many whole plant models is currently represented solely by the accumulation of biomass components. The introduction of GrOE-FBA-driven expanding cells could help improve the representation of metabolism in these models, potentially improving their predictive power.

## MATERIALS AND METHODS

### Plant material and growth conditions

Tomato (*Solanum lycopersicum* cv. “Moneymaker”) seeds were sterilized and germinated on Murashige-Skoog (MS) medium. The plants were grown in a glasshouse in a 16 h photoperiod at 22 to 23°C day temperature and 20 to 22°C night temperature with supplementary lighting to maintain an irradiance of 250 to 400 μmol m^-2^ s^-1^. Lateral stems were systematically removed. Each flower anthesis was recorded, and trusses were pruned at five developed fruits to limit fruit size heterogeneity. Tomato fruits were harvested at nine different developmental stages corresponding to 8, 15, 22, 28, 34, 42, 50, 52 and 59 DPA with the last four corresponding to mature green, turning, orange and red fruit developmental stages respectively. Fruit samples were collected from the first two levels of trusses in the plant. All materials were frozen in liquid nitrogen and storage at −80°C until use.

### Fruit biomass measurements and morphology analysis

Cell wall was extracted according to an established protocol (Ruprecht *et al.*, 2011). The remaining insoluble material was washed with water and ethanol, air-dried and weighed. Lipids were extracted from a known mass of ground tissue using the chloroform/methanol protocol (Bligh and Dyer, 1959). Protein extracted with 6 M urea, 2 M thiourea buffer (Salem *et al.*, 2016) was quantified using the Bradford assay (Bradford, 1976). The amino acid content of protein hydrolysates (50% w/v trichloroacetic acid, 6 M HCl, 100°C/24 h) (Antoniewicz *et al.*, 2007) was determined by an established GC/EI-TOF-MS protocol (Luedemann *et al.*, 2008; Fernie *et al.*, 2011; Osorio *et al.*, 2011). Starch content was determined by enzymatic digestion and spectrophotometric assay of the resultant glucose (Hendriks *et al.*, 2003). Fruit height and diameter were measured using a calibrated digital Vernier. The fresh weight to dry weight ratio (Fw/Dw) was determined by weighing fresh tissues and reweighing them after drying to constant weight in a forced-air oven at 65°C.

### Harvesting phloem exudates

Exudates were harvested from tissues corresponding to mature green and red fruit pedicels using the EDTA-facilitated method (Tetyuk *et al.*, 2013).

### Gas-chromatography mass spectrometry of fruit content

Metabolite analysis of fruit samples was performed using gas chromatography coupled to electron impact ionization-time of flight-mass spectrometry (GC/EI-TOF-MS) (Lisec *et al.*, 2006). Plant material was extracted using the method described elsewhere (Osorio *et al.*, 2011). Data analysis was performed using ChromaTOF 1.0 (Leco, www.leco.com) and TagFinder v.4.0 (Luedemann *et al.*, 2008). Cross-referencing of mass spectra was performed with the Golm Metabolome database (Kopka *et al.*, 2005). Documentation of metabolite profiling data acquisition is reported following recommended guidelines (Fernie *et al.*, 2011). Curve fitting (Colombié *et al.*, 2015) was used to generate curves for metabolite content and fluxes based on experimental measurements.

### Flux balance analysis and flux variability analysis

Metabolic reactions associated with phospholipid biosynthesis (PC, PE and PA), amino acid catabolism, nucleotide biosynthesis, β oxidation, lycopene biosynthesis and phytol metabolism were added to a previously published mass and charge-balanced model of primary metabolism in plant cells (Shameer *et al.*, 2018) to generate the model PlantCoreMetabolism_v1_2. A complete log of all model curations and associated literature is presented in Table S2. Parsimonious FBA (pFBA) and FVA functions available in cobrapy (Ebrahim *et al.*, 2013) version 0.13.4 were updated to perform weighted-pFBA and weighted-FVA, respectively. Aggregator reactions used to measure fruit metabolite content were given a zero weight while all other reactions were given a weight of 1. Cobrapy in Jupyter notebook (python 2.7) was used to run and document all scripts (https://github.com/ljs1002/Shameer-et-al-Predicting-metabolism-during-growth-by-osmotic-cell-expansion).

### Estimating maintenance costs

Non-growth associated maintenance (NGAM) costs in all models were represented by flux through an ATPase and three NADPH oxidase (cytosolic, mitochondrial and plastidic) pseudo-reactions, constrained to a 3:1 (ATP hydrolysiss: NADPH oxidase) ratio based on the results of a previously published study (Cheung *et al.*, 2013).

Dividing cells were assumed to have a carbon conversion efficiency (CCE) of 70% based on previously published data (Chen and Shachar-Hill, 2012). Fluxes through the NGAM reactions in the dividing cell model were gradually increased until a CCE~70% was achieved. When the NGAM ATPase flux was constrained to 0.062 mmol/mL/day, the CCE of the system was observed to be 69.92%. This NGAM ATPase flux was then used to constrain maintenance in the dividing cell model. The same NGAM constraints were imposed on the expanding cell model.

Each phase of the multiphase GrOE-FBA model was 5.1 days long. The mean respiratory cost of NGAM based on data reported by Walker and Thornley (1977) was assumed to be 1.7 mmol CO2/fruit/day throughout fruit development. This is equivalent to 8.5 mmol CO2/fruit/5.1 days. Using a model of primary metabolism in pericarp cell constrained so as to prevent the net synthesis of any metabolite, an NGAM ATPase flux of 26.2 mmol/fruit/5.1 days was found to result in a respiration rate of 8.5 mmol CO2/fruit/5.1 days. This NGAM ATPase flux was used to constrain NGAM in all phases of fruit development.

### Modelling primary metabolism in a dividing heterotrophic cell suspension

The PlantCoreMetabolism_v1_2 model, was used to generate representations of metabolism in dividing tomato cells. Previously published data from tomato heterotrophic cell suspension cultures report a maximum growth rate of 2 mg DW/mL/day (Rontein *et al.*, 2002). Data on biomass content (cell wall, sugars, organic acids and protein) of cells demonstrating the maximum growth rate, from the same study, was used to generate a biomass equation for the dividing cell system as in conventional FBA modelling. The lipid: cellulose content ratio reported in Arabidopsis cell cultures (Williams *et al.*, 2010) were used to estimate the lipid content of the cells and the biomass equation was updated accordingly. The biomass accumulation rate of the model was constrained to 2 mg DW/mL/day. For the sake of simplification, the cell wall and cell membrane were assumed to be composed of only cellulose and phosphatidic acid (PA) respectively. Constraints based on equation 2 were introduced to promote solute partitioning between cytosol and vacuole. Because of the lack of data available in heterotrophic tomato cells, the value for Vv/Vc required for equation 2 was calculated from data published on tomato pericarp subcellular volume fractions (Beauvoit *et al.*, 2014). NGAM was represented in the model as described earlier. Glucose was set as the sole carbon source and FBA with minimization of sum of fluxes as the objective was used to predict the optimal flux distribution.

### Modelling primary metabolism in a system of expanding 26 DPA pericarp cells

The PlantCoreMetabolism_v1_2 model, was used to generate representations of metabolism in expanding 26 DPA tomato pericarp cells. The rate of cell expansion was estimated to be at its highest in 26 DPA tomato pericarp cells (0.174 nL/cell/day; estimated from data in Beavuoit et al., 2014). For the sake of consistency, the number of cells in the expanding cell system was kept equal to that in the dividing cell suspension when growth rate attained the maximum rate of 2 mg DW/mL/day (1.45×10^6^ cells/mL). The composition of soluble metabolites in 26 DPA pericarp cells were estimated from third-degree polynomial curves fitted to published experimental data (Colombié *et al.*, 2015). Equations 1 and 2 were used to introduce osmotic constraints in the model and the values for *V_cell_* and *C_cell_* were estimated from published data (Almeida and Huber, 1999; Beauvoit *et al.*, 2014). The difference in osmotic content between DPA 26 and 27 was used to set the demand for osmolytes. Data on subcellular volume fractions published by Beauvoit *et al.* were used to calculate V_v_/V_c_ at 26 DPA. The difference between cellulose, lipid and protein content between DPA 26 and DPA 27, as calculated from equation 3 - 5, was used to set the demand for these biomass elements. NGAM was was represented in the model as described earlier. Glucose was set as the sole carbon source (as in the dividing-cells model) and FBA was used to predict metabolic fluxes that minimized the sum of fluxes while maximizing the organic content.

### Implementation of GrOE-FBA constraints in the multiphase developing fruit model

Tomato fruit development from 13.1 to 43.7 DPA (mature green fruits observed at 42 DPA) is mainly due to expansion of existing cells (Gillaspy *et al.*, 1993) and GrOE-FBA was used to model metabolism during these stages. Assuming the tomato fruit is composed of uniformly packed pericarp cells, equation 1 cam be transformed to the following form for a whole fruit,

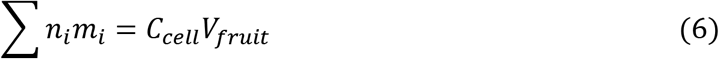

where *i* is a metabolite/ion in the fruit and *V_fruit_* is the volume of the fruit.

Constraints based on equation 2 and 6 were imposed on each phase of the multiphase developing fruit model. Previously published data on subcellular volume fractions (Beauvoit *et al.*, 2014) were used to calculate *V_v_/V_c_* during the different stages of fruit development and *C_cell_* was estimated based on osmolalities reported in the literature (Almeida and Huber, 1999). Values for *V_fruit_* for all 10 phases of fruit development modelled were determined experimentally (Table S1).

Pericarp cell volume for each phase of the multiphase fruit model was calculated from published data (Beauvoit *et al.*, 2014), and cellulose, phospholipid and protein demand fluxes were imposed on the system using equations 3, 4 and 5 and scaling them from μmol/cell to mmol/fruit.

Carbon influx rate into developing tomato fruits has been reported to be inversely proportional to fruit carbon content and size (Walker and Ho, 1977a,b). A hyperbolic curve was fitted to capture the relationship between carbon influx rates and fruit carbon content more accurately. Fruit carbon content has been reported to be linearly related to fruit volume (Walker and Ho, 1997b). Hence, initial carbon content for each developmental phase was calculated based on fruit volume at the respective phases. These values were then used to predict the upper bounds for the flux of nutrients from the phloem in the respective phases of the developing tomato model.

