## Appendix S1 for "Predicting metabolism during growth by osmotic cell expansion"

APPENDIX SI: Derivations for equations 1-5

**Derivation of equation 1**

The osmolarity of a cell ($C_{cell}$) can be described by the following equation,

$$C_{cell}= \sum\varphi_{i}n_{i}c_{i}$$

where $\varphi_{i}$, $n_{i}$ and $c_{i}$ are the osmotic coefficient of the solvent, the van’t Hoff factor and molar concentration of molecule *i* in the cell (Goodman, 2008).

The osmotic coefficient is a property of the solution that characterizes its deviation from ideal behaviour stated by Raoult’s law. The van’t Hoff factor of a solute is the ratio between the number of particles formed when a solute is dissolved in a solution and number of moles of solute dissolved.

For an ideal solution, the osmotic coefficient is 1, the van’t Hoff factor of a non-ionic solute is 1, and the van’t Hoff of an ionic solute is equal to the number of ions when the solute is completely disassociated. For example, in an ideal solution, the van’t Hoff factor of glucose, NaCl and CaCl_2_ are 1, 2 and 3 respectively.

Assuming the intracellular medium behaves as an ideal solution,

$$\begin{aligned} C_{cell}= \sum n_{i}c_{i} \#(A) \end{aligned}$$

Replacing concentration ($c_{i}$) with number of moles ($m_{i}$) and volume of the cell ($V_{cell}$) gives

$$C_{cell}= \sum n_{i}\frac{m_{i}}{V_{cell}}$$

$$\begin{aligned} \sum n_{i}m_{i}=C_{cell}V_{cell} \#\left( 1 \right) \end{aligned}$$

**Derivation of equation 2**

Vacuolar expansion is the result of osmotic flux ($J_{w}$) into the vacuole, which can be described by

$$\begin{aligned} J_{w}= A_{v}\left( \Psi_{c}-\Psi_{v} \right)\#\left( B \right) \end{aligned}$$

where $A_{v}$ is the hydraulic conductivity of the tonoplast, $\Psi_{c}$ is the cytosolic water potential and $\Psi_{v}$ is the vacuolar water potential. Water potential ($\Psi$) can be expressed in terms of hydrostatic or turgor pressure ($P$) and osmotic pressure ($\Pi$) (Passioura, 1982)

$\begin{aligned} \Psi=P- \Pi\end{aligned}$

Replacing values of $\Psi_{c}$ and $\Psi_{v}$ in equation B gives

$J_{w}= A_{v}(P_{c}-\Pi_{c}-P_{v}+\Pi_{v})$

and since $P_{c}$ = $P_{v}$

$J_{w}= A_{v}(\Pi_{v}-\Pi_{c})$

At steady state $J_{w}$ = 0 and since $A_{v}$ ≠ 0,

$\begin{aligned} \Pi_{v}= \Pi_{c} \#\left( C \right) \end{aligned}$

From the van’t Hoff relation,

$$\Pi=RTC$$

where R is the gas constant and T is the absolute temperature. Hence equation C can be expressed in terms of the cytosolic and vacuolar solute concentrations

$$C_{v}= C_{c}$$

and using equation A gives,

$$\begin{aligned} \sum n_{j}c_{j}=\sum n_{i}c_{i} \#\left( D \right) \end{aligned}$$

where $n_{a}$ and $c_{a}$ are the van’t Hoff factor and molar concentration of a metabolite $a$ respectively; and $i$ and $j$ are metabolites in the cytosol and vacuole respectively. Rewriting the molar concentrations in terms of the number of moles of the metabolite *j* ($m_{j}$) and the volume of the vacuole ($V_{v}$) gives,

$$c_{j}=\frac{m_{j}}{V_{v}}$$

Similarly, for the cytosol,

$$c_{i}=\frac{m_{i}}{V_{c}}$$

where $m_{i}$ and $V_{c}$ are the number of moles of metabolite $i$ in cytosol and the volume of the cytosol respectively. Using the values of $c_{j}$ and $c_{i}$ in equation D, leads to

$$\sum n_{j}\frac{m_{j}}{V_{v}}=\sum n_{i}\frac{m_{i}}{V_{c}}$$

$\frac{\sum n_{j}m_{j}}{\sum n_{i}m_{i}}=\frac{V_{v}}{V_{c}}$ $\left( 2 \right)$

**Derivation of equations 3 and 4**

In order to estimate the demand for cellulose, phospholipids and protein as a result of cell expansion, a simple representation of the plant cell was created by assuming that: (a) cells are cube-shaped; (b) the cell wall is uniformly thick and composed of only cellulose; and (c) the cell membrane is uniformly thick. If $a$ is the edge of the cube-shaped cell at time $t$, $b$ is the thickness of the cell wall and $c$ is the thickness of the cell membrane (as in Figure 2b), then the volume of cell wall at $t$ can be represented by

$$V_{cellwall,t}=V_{totalcell,t}- V_{cellwithoutcellwall,t}$$

where $V_{totalcell,t}$ and $V_{cellwithoutcellwall,t}$ are the volumes of the entire cell and of the cell without its cell wall respectively, at time $t$. This can be expressed in terms of $a$ and $b$ as

$$V_{cellwall,t}=a^{3}- \left( a-2b \right)^{3}= 8b^{3}+6a^{2}b-12ab^{2}$$

$$V_{cellwall,t}= 8b^{3}+6{{b(V}_{totalcell,t})}^{\frac{2}{3}}-12b^{2}({V_{totalcell,t})}^{\frac{1}{3}}$$

Since the cell wall is assumed to be composed of only cellulose, $V_{cellwall,t}= V_{cellulose,t}$, and the cellulose content of the cell in grams (${mass}_{cellulose,t}$) at time $t$ is given by

$${mass}_{cellulose,t}= V_{cellwall,t}\rho_{cellulose}$$

$$=\left\{ 8b^{3}+6{{b(V}_{totalcell,t})}^{\frac{2}{3}}-12b^{2}({V_{totalcell,t})}^{\frac{1}{3}} \right\}\rho_{cellulose}$$

where $\rho_{cellulose}$ is the density of cellulose. Then, since cellulose is represented in the model in glucose units, the cellulose content expressed in moles for the model at time $t$ (${moles}_{cellulose,t}$) can be calculated using the molecular weight of glucose (${MW}_{glucose}$):

$$\begin{aligned} {moles}_{cellulose,t}= \frac{8b^{3}+6{{b(V}_{totalcell,t})}^{\frac{2}{3}}-12b^{2}({V_{totalcell,t})}^{\frac{1}{3}}}{{MW}_{glucose}}\rho_{cellulose}\#(3) \end{aligned}$$

Equation 3 can be used to estimate the cellulose content of an expanding cell.

Similarly, the cell membrane phospholipid content of a cube-shaped cell at time $t$ can be calculated from the simple cell representation (Figure 2b) as follows

$$V_{cellmembrane,t}=V_{totalcell,t}- V_{cellwall,t}- V_{cytoplasm,t}$$

where $V_{cellmembrane,t}$ and $V_{cytoplasm,t}$ are the volumes of cell membrane and cytoplasm respectively

$$V_{cellmembrane,t}= 8c^{3}+6a^{2}c-24b^{2}c-12c^{2}\left( a-2b \right)-24abc$$

$$V_{cellmembrane,t}=8c^{3}+6c({V_{totalcell,t})}^{\frac{2}{3}}-24b^{2}c-12c^{2}\left\{ ({V_{totalcell,t})}^{\frac{1}{3}}-2b \right\}-24bc({V_{totalcell,t})}^{\frac{1}{3}}$$

The amount of phospholipid (in moles) in a cell at time $t$ (${moles}_{phospholipid,t}$) is

$${moles}_{phospholipid,t}=\frac{{mass}_{phospholipid,t}}{{MW}_{phospholipid}}= \frac{V_{cellmembrane,t}}{{MW}_{phospholipid}}\rho_{phospholipid}$$

where ${mass}_{phospholipid,t}$ is the amount of phospholipid (in grams), $\rho_{phospholipid}$ is the density of membrane phospholipids, ${MW}_{phospholipid}$ is the molecular weight of membrane phospholipids. Using the value of $V_{cellmembrane,t}$ derived from Figure 2b, gives

$$\begin{aligned} {moles}_{phospholipid,t}= \\ \frac{8c^{3}+6c({V_{totalcell,t})}^{\frac{2}{3}}+24b^{2}c-12\left\{ ({V_{totalcell,t})}^{\frac{1}{3}}-2b \right\}-24bc({V_{totalcell,t})}^{\frac{1}{3}}}{{MW}_{phospholipid}}\rho_{phospholipid} \end{aligned}$$

This equation can be used to estimate the cell membrane associated phospholipid content of a cell. However, only a fraction of the total cell phospholipid content is associated with cell membrane ($f_{PM}$), and so the equation is modified,

$$\begin{aligned} {moles}_{phospholipid}= \\ \frac{8c^{3}+6c({V_{totalcell})}^{\frac{2}{3}}+24b^{2}c-12\left\{ ({V_{totalcell})}^{\frac{1}{3}}-2b \right\}-24bc({V_{totalcell})}^{\frac{1}{3}}}{f_{PM}{MW}_{phospholipid}}\rho_{phospholipid} \#(4) \end{aligned}$$

Equation 4 can be used to estimate the phospholipid content of an expanding cell.

**Derivation of equation 5**

In order to determine the protein content of a cell based on its volume, two assumptions were made. First, the fraction of the total protein content localized to the vacuole was assumed to be negligible. Secondly, a cell during expansion was assumed to maintain the protein concentration in the cytoplasm to maintain functionality. With these assumptions, the total protein content of the cell can be estimated as follows

$$\begin{aligned} {moles}_{protein,t}=C_{protein}V_{cytoplasmic,t} \#(5) \end{aligned}$$

where ${moles}_{protein,t}$ is the amount of protein (in mols) in a cell at $t$, $C_{protein}$ is the molar concentration of protein in the cytoplasm and $V_{cytoplasmic,t}$ is the volume of the cytoplasm at time $t$.

Equation 5 can be used to estimate the protein content of an expanding cell.
