## Supplementary material for "Predicting metabolism during growth by osmotic cell expansion": Figure S1

**Supplemental Figure S1: Comparison of predicted and measured metabolite contents in developing tomato fruit**

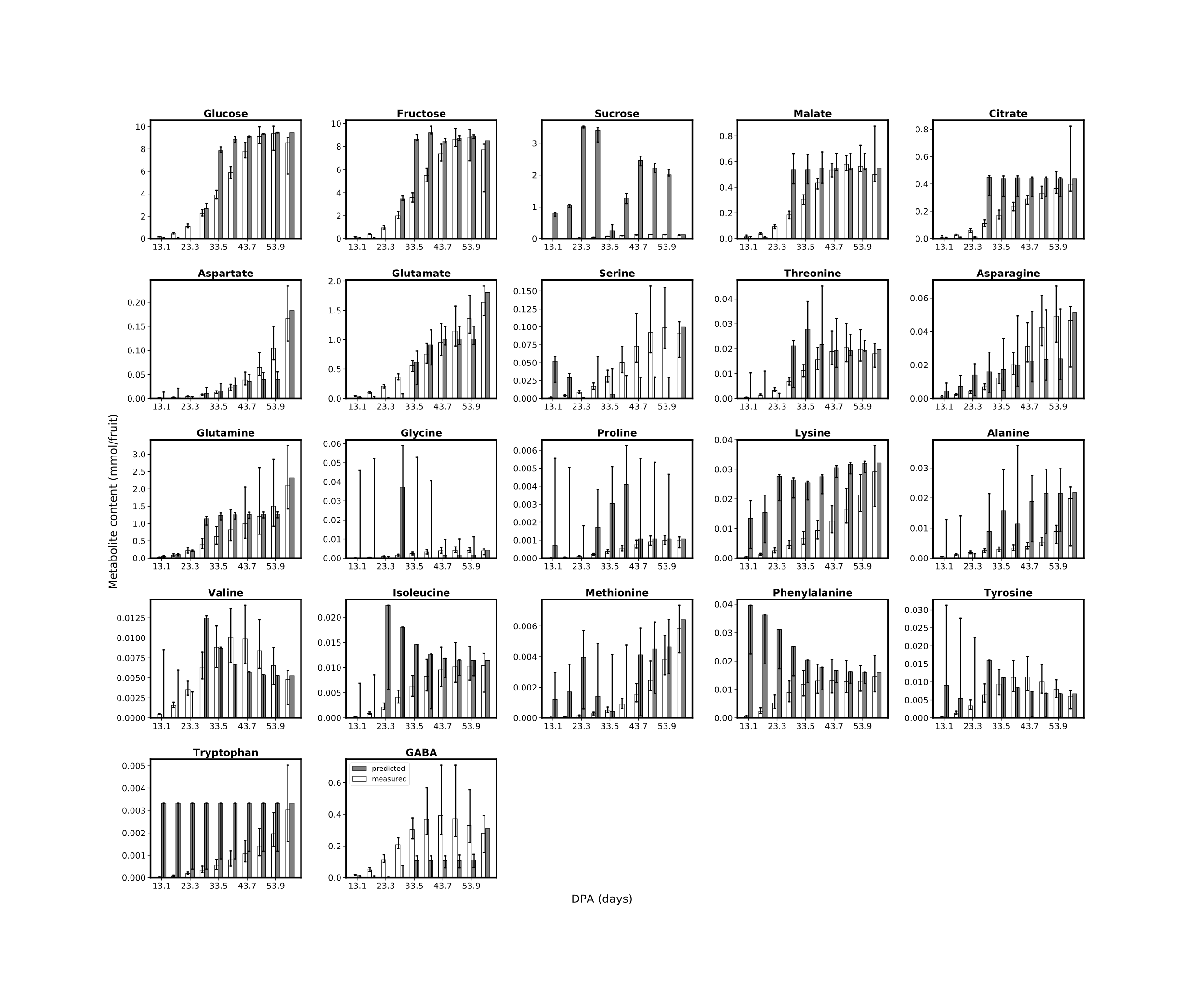
