## Supplementary material for "Predicting metabolism during growth by osmotic cell expansion": Table S1

**Supplemental Table S1: List of parameters used in the study.**

| **Parameter** | **Value** | **Source** |
| --- | --- | --- |
| cell wall thickness (b) | 100 nm | Alberts *et al.*, 2008 |
| cell membrane thickness (c) | 4 nm | Milo *et al.*, 2009 |
| density of 1,2-dipalmitoyl-sn-glycero-3-phosphatidylcholine ($\rho_{PC}$) | 1 g/m^3^ | Kupiainen *et al.*, 2005 |
| van der Waal’s volume of 1,2-dipalmitoyl-sn-glycero-3- phosphatidic acid | 698.93 Å^3^ | Cotter *et al.*, 2006 |
| density of 1,2-dipalmitoyl-sn-glycero-3-phosphatidylethanolamine ($\rho_{PE}$) | 1.01 g/m^3^ | https://www.scbt.com/scbt/product/1-2-dipalmitoyl-sn-glycero-3-phosphoethanolamine-923-61-5 |
| **Tomato fruit** |  |  |
| cell volume ($V_{totalcell}$) | $\frac{0.0289433}{0.00760074+ e^{(-0.18324543DPA)}}-0.03277816$ m^3^ | Beauvoit *et al.*, 2014 |
| vacuole volume fraction ($V_{v}$) | $0.853\left( 1-e^{\left( \frac{-2292-t^{*}}{10633} \right)} \right)$ | Beauvoit *et al.*, 2014 |
| cytoplasm volume fraction ($V_{c}$) | $\frac{0.933- V_{v}}{1.13}$ | Beauvoit *et al.*, 2014 |
| cell protein concentration | 19.66 mol/m^3^ | Experimentally determined |
| fruit volume ($V_{fruit}$) | $\frac{-0.000085358704}{1+ \left( \frac{{DPA}^{**}}{28.56023} \right)^{9.692893}}+ 0.0000889527$ m^3^ | Experimentally determined |
| *$t$ is time in minutes  **$DPA$ stands for days post anthesis | | |

**Alberts, B., Jonhson, A., Lewis, J., Morgan, D. and Raff, M.** (2008) Cell junctions and the extracellular matrix. In *Molecular Biology of the Cell*. Taylor and Francis, pp. 1035–1090.

**Beauvoit, B.P., Colombié, S., Monier, A., et al.** (2014) Model-assisted analysis of sugar metabolism throughout tomato fruit development reveals enzyme and carrier properties in relation to vacuole expansion. *Plant Cell* **26**, 3224–3242. http://doi.org/10.1105/tpc.114.127761

**Cotter, D., Maer, A., Guda, C., Saunders, B. and Subramaniam, S.** (2006) LMPD: LIPID MAPS proteome database. *Nucleic Acids Res.* **34**, D507-10. https://doi.org/10.1093/nar/gkj122

**Kupiainen, M., Falck, E., Ollila, S., Niemelä, P., Gurtovenko, A.A., Hyvönen, M.T., Patra, M., Karttunen, M. and Vattulainen, I.** (2005) Free volume properties of sphingomyelin, DMPC, DPPC, and PLPC bilayers. *J. Comput. Theor. Nanosci.* **2**, 401–413. http://doi.org/10.1166/jctn.2005.211

**Milo, R., Jorgensen, P., Moran, U., Weber, G. and Springer, M.** (2009) BioNumbers The database of key numbers in molecular and cell biology. *Nucleic Acids Res.* **38**, 750–753. http://doi.org/10.1093/nar/gkp889
